# Genomic insights into polysaccharide substrate utilization and novel genera resource mining in macroalgal epiphytic bacteria

**DOI:** 10.1101/2025.04.07.647558

**Authors:** Tian-He Liu, Hao-Yu Zhou, Han-Zhe Zhang, Feng-Qing Wang, Ye-Zi Liu, Jin-Hao Teng, Si-Jia Fan, De-Chen Lu, Zong-Jun Du

## Abstract

Marine macroalgae, among the fastest photosynthesizing organisms, play a crucial role in the ocean carbon cycle by converting fixed carbon dioxide into polysaccharides. Macroalgal epiphytic bacteria possessing specific polysaccharide utilization loci (PULs) have a generalized polysaccharide degradation potential that facilitates their growth and colonization. In this study, we conducted extensive research on their polysaccharide degradation potential. Through sampled and purified epiphytic bacteria and metagenomic analysis revealed a high prevalence of novel genera and species. Two novel genera, 1117^T^ and 3-347^T^, were taxonomically characterized and conducted detailed functional analyses. The results demonstrated that these genera possess abundant PULs and strong capabilities for synthesizing secondary metabolites. Furthermore, their high relative abundance on macroalgal surfaces aligns with global ecological distribution patterns. These traits facilitate their colonization, growth, and environmental adaptation on macroalgal surfaces. We further performed in-depth annotation of a large number of PULs and CAZyme genes of macroalgal epiphytic bacteria. Potential polysaccharide substrates for their degradation can be predicted and focused on. Additionally, we conducted growth curve analyses by starch, xylan, β-1,3-glucan, and carboxymethyl cellulose substrates to validate the genomic predictions. In summary, our findings demonstrate that macroalgal epiphytic bacteria possess significant potential for degrading algal polysaccharides. This capability may enhance their competitiveness and survival probability on macroalgal surfaces. These bacteria, originating from different sources and genera, possess similar PULs, which may result from horizontal gene transfer or evolutionary relationships.

**Importance:** Macroalgae are major primary producers in coastal areas and their carbon sequestration capacity per unit area far exceeds that of terrestrial forests. In this work, we extensively studied macroalgal epiphytic bacteria with polysaccharide degradation potential. We found that epiphytic bacteria from different macroalgal sources and genera share similar PULs to degrade the same polysaccharide, which may be the result of horizontal gene transfer or evolutionary relationships. Core taxa on the macroalgal surface have gradually evolved polysaccharide-degrading abilities of different marine macroalgae in order to expand their colonization and survival chances. We also identified a large number of uncultivated algal biosphere species and unreported new genera and species for expansion of macroalgal epiphytic bacteria studies. These studies have thus highlighted the important ecological and research value of macroalgal epiphytic bacteria, especially in influencing polysaccharide carbon storage and marine carbon cycling.

## Introduction

Macroalgae are photosynthetic, multicellular eukaryotic organisms (1) that are major primary producers in coastal areas and global carbon sinks (2, 3). Carbon sequestration capacity per unit area of macroalgae far exceeds that of terrestrial forests (3). Macroalgae contain a variety of highly sulfated polysaccharides, most notably fucoidan, ulvan and carrageenan, which form key components of carbon fixation by macroalgae (4–6). The surface of the macroalgae is colonized by bacteria called macroalgal epiphytic bacteria, both of which have co-evolved for roughly 1.6 billion years in a complex relationship (7). They interact and influence each other, playing a crucial role in the ecological and biochemical processes of the coastal ocean (8). Polysaccharides, which are macromolecular polymers composed of more than 10 monosaccharides with glycosidic linkages, and are the main bioactive component of macroalgae (9). These complex polysaccharides are abundant in the extracellular matrix (ECM) of macroalgae and constitute a significant portion of their dry weight (10). Through photosynthesis, marine macroalgae sequester large amounts of carbon dioxide into polysaccharides, which account for approximately 50% of macroalgal biomass (11). As a central metabolic fuel in the marine carbon cycle (12), polysaccharides serve critical functions as energy storage molecules, cell wall components, and extracellular defense mechanisms (13). In natural environments, the degradation and cycling of polysaccharides are primarily regulated by microorganisms, with the biocatalytic decomposition of polysaccharides being accomplished through the combined action of microbial communities (14). Among these bacteria, *Bacteroidota* are the dominant phylum and have been reported to possess a high capacity for polysaccharide utilization (15). For degradation of macroalgal polysaccharides, bacteria occupying the epiphytic ecological niche of macroalgae are most likely to be the main degraders of carbon fixed by marine macroalgae (16, 17). However, the role of macroalgae has long been ignored in the discussion of the ocean carbon cycle until recent years when it has received more attention (18). Consequently, marine macroalgae and their epiphytic bacteria are integral to the ecological and biochemical dynamics of coastal oceans.

The phylum *Bacteroidota* is widely recognized as a major utilizer of high-molecular-mass dissolved organic matter in marine ecosystems (19). Its members play a crucial role as heterotrophs, contributing significantly to the cycling of organic carbon in aquatic habitats (20). These bacteria are particularly adept at targeting algal polysaccharides, with the genes responsible for the breakdown and uptake of polysaccharides often co-located in specialize PULs (21). Beyond marine environments, *Bacteroidota* are also key polysaccharide degraders in terrestrial ecosystems, notably in the human gut (13). This dual role underscores their importance in both aquatic and terrestrial carbon cycling.

The PULs were first reported by Tancula et al. (22) and was an important component of the unique biological mechanism of polysaccharide utilization in the phylum *Bacteroidota.* Polysaccharide degradation in *Bacteroidota* was usually encoded in specialized genomic islands that are referred to as PULs (23). PULs contained a gene tandem coding for a SusD-like glycan-binding protein and a SusC-like/ TonB-dependent transporter as well as dedicated CAZymes and accessory proteins such as sulfatases to orchestrate binding, uptake and degradation of a particular polysaccharide (24). In *Bacteroidota*, CAZymes usually consisted of clusters of co-regulated genes involved in carbohydrate binding, hydrolysis and transport used to catalyze the direct degradation of polysaccharides (25). CAZymes included glycoside hydrolase (GH), glycosyltransferase (GT), polysaccharide lyase (PL), carbohydrate esterase (CE), auxiliary activities (AA) and carbohydrate binding module (CBM). PUL of the *Bacteroidota* produce specific enzymes depending on the complex polysaccharide to be degraded. Conversely, the composition of the enzymes produced by PUL can provide information about the structure of the targeted polysaccharide (26). Therefore, the genetic composition of PUL was used to predict the polysaccharide niche of a bacterium (27).

Here, we investigated the polysaccharide degradation potential of macroalgal epiphytic bacteria (including strains 1117^T^ and 3-347^T^). We conducted in-depth annotation of a large number of as-yet-undescribed PULs and diverse CAZyme genes within the epiphytic microbial community. By analyzing the widespread distribution of PULs and the diversity of CAZymes, we predicted potential polysaccharide substrates targeted by these bacteria. Metagenomic analysis revealed that epiphytic bacteria from different macroalgal sources and genera share similar PULs when degrading the same polysaccharides, highlighting a conserved functional mechanism. We further identified and analyzed key polysaccharide substrates targeted by macroalgal epiphytic bacteria and validated their degradation potential through growth curve experiments using selected substrates.

## Results

### Isolation, cultivation, analysis and identification of macroalgal epiphytic bacteria

We sampled three different marine macroalgae (red, green and brown macroalgae), surrounding seawater and surface sediments in a coastal area of Weihai, China. The macroalgae samples included *Ulva* sp., *Saccharina* sp., *Grateloupia* sp. and *Gelidium* sp. Bacteria were isolated from the above samples by extraction and dilution, cultured and purified with culture media (Methods for extraction, dilution, isolation, cultivation, and purification of bacteria are described in Lu et al.) (28). After 21 days of incubation at 28.0 °C, abundant single colonies were observed, predominantly composed of *Bacteroidota*.

We analyzed the relative abundance of *Bacteroidota* across different macroalgae samples (Fig. 1; Table S1). The relative abundance of *Bacteroidota* and its mean value were highest in *Ulva* sp., followed by *Grateloupia* sp. and *Gelidium* sp., the lowest in *Saccharina* sp. Notably, phylum *Bacteroidota* exhibited the highest relative abundance in our samples, suggesting that macroalgae-colonizing *Bacteroidota*. Additionally, we quantified the number of novel species and genera within the phylum *Bacteroidota* and calculated their respective percentages in different macroalgae samples (Fig. 1; Table S1). The analysis revealed that *Grateloupia* sp. harbored the highest proportion of novel species and genera, whereas *Saccharina* sp. contained the lowest. However, in terms of the mean proportion of novel genera, *Saccharina* sp. exceeded that of *Ulva* sp. These findings demonstrate that *Bacteroidota* represents a core taxon on the surfaces of the macroalgae sampled, with a significant presence of novel genera and species. This suggests that macroalgal surfaces are enriched with bacteria, highlighting their considerable research potential for the discovery and characterization of novel macroalgal epiphytic bacterial species.

**Fig. 1.**
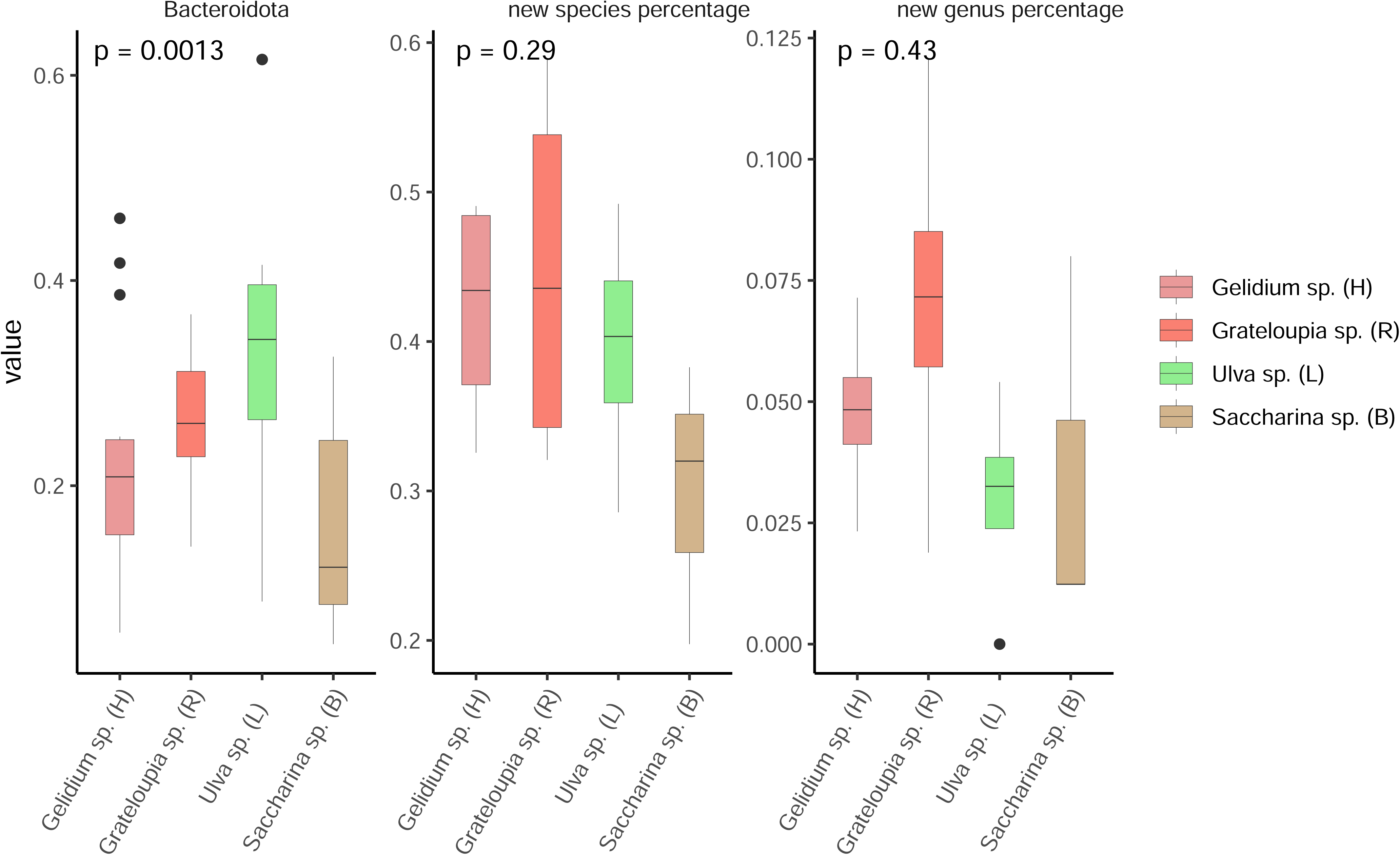
We analyzed the relative abundance of *Bacteroidota* in different kinds of macroalgae samples. The result shows that the relative abundance and the average value of the relative abundance of *Bacteroidota* was highest in *Ulva* sp., followed by *Grateloupia* sp. and *Gelidium* sp., the lowest in *Saccharina* sp. Then, we counted the number of new species and genera within the phylum *Bacteroidota* and analyzed the percentages of them in four different macroalgae samples. According to the box plot, we found that the highest percentages of new species and genera in samples of *Grateloupia* sp. and lowest in samples of *Saccharina* sp. However, for average value of the percentage of new genera, the *Saccharina* sp. is higher than *Ulva* sp., which is different with the result in the box plot.

In order to expand the macroalgal epiphytic bacteria resources and to study their potential capabilities and ecological significance, we identified two new genera of *Bacteroidota* isolated from green macroalgae (1117^T^) and red macroalgae (3-347^T^), respectively (Fig. S1B; S1C). We used multiphase taxonomy to validate their taxonomic status, performed detailed functional genomics studies and extensive experimental validation. These results suggest that they represent novel genera within their respective taxonomic groups and have great research potential.

The complete 16S rRNA gene sequence of strain 1117^T^ (1,521 bp) and 3-347^T^ (1,524 bp) was obtained from the genome. Strain 1117^T^ exhibited the highest 16S rRNA gene sequence similarity to *Cellulophaga tyrosinoxydans* EM41^T^ (93.9 %), while the highest similarity to 3-347^T^ was *Maribacter vaceletii* DSM 25230^T^ (93.3 %). According to Yarza et al. (29), a 16S rRNA gene sequence identity less than or equal to 94.5 % strongly supports the distinction of different genera. Phylogenetic analysis of MEGA revealed that they formed distinct branches within the family *Flavobacteriaceae* (Fig. S1A), representing two novel genera, *Ulvaetocola* and *Agarflavus*. Consistent topologies were observed with ML and ME algorithms (Fig. S2).

The draft genome sequencing showed that strain 1117^T^ had a genome length of 4,292,054 bp with 36.6 mol% DNA G+C content and 3-347^T^ had a genome length of 4,342,517 bp with 35.9 mol% G+C content. Detailed genome information of strains 1117^T^, 3-347^T^ and all bacteria in the 16s rRNA phylogenetic tree were shown in Table S2. The POCP and AAI values between strain 1117^T^ and its most closely related species were 53.5 %-60.7 % and 70.3 %-71.6 %, respectively. Similarly, the POCP and AAI value between strain 3-347^T^ and *Maribacter vaceletii* DSM 25230^T^ was calculated to be 55.8 % and 70.1 %, respectively. Further, we constructed the genomic phylogenetic tree and compare AAI and POCP values including all available genomes from the 16S rRNA phylogenetic tree, four macroalgal epiphytic bacteria and two *Ulvaetocola* metagenome-assembled genomes (MAGs) (Fig. S3). These values consistently fell within the established genus delineation thresholds (Table S3) (POCP >50.0%, AAI >65.0%) (30, 31). Additionally, all ANI and dDDH values (Table S4) in the genomic phylogenetic tree were well below species delineation thresholds (ANI 95.0–96.0%, dDDH 70%) (32, 33).

We also determined their taxonomic status through comprehensive phenotypic features, physiological and chemotaxonomic characteristics (detailed information in Supplementary Materials, Fig. S4; Tables S5, S6). And Table S7 summarized the characteristics of the twenty-three species of *Flavobacteriaceae* most closely related to strains 1117^T^ and 3-347^T^, highlighting key features that were carefully organized. The reference strain for physiological and chemotaxonomic comparisons of strain 1117^T^ was *Cellulophaga tyrosinoxydans* EM41^T^, and that of 3-347^T^ was *Maribacter vaceletii* DSM 25230^T^. Hence, we proposed strains 1117^T^ and 3-347^T^ represented two novel genera isolated from macroalgae surface of the phyla *Bacteroidota*, for which the names *Ulvaetocola algae* gen. nov., sp. nov. and *Agarflavus algae* gen. nov., sp. nov. were proposed with the type strains 1117^T^ (=MCCC 1H00987^T^= KCTC 102106^T^) and 3-347^T^ (=MCCC 1H00853^T^= KCTC 102101^T^), respectively.

## Genomic Function Analysis

### Prediction of Secondary Metabolites

Analyzing the potential functions of all macroalgal epiphytic bacteria was beyond the scope of our study, so we selected two new genera for detailed genomic function analysis. Firstly, we analyzed the secondary metabolite biosynthesis gene clusters (BGCs) of strains 1117^T^ and 3-347^T^ (Fig. S5). Strain 1117^T^ contained fifteen putative BGCs, while strain 3-347^T^ contained seven putative BGCs. Both strains shared BGCs encoding the production of terpenes, NRPS and Type III Polyketide synthase (Type III PKS). The most abundant types of BGCs in 1117^T^ encoded for non-ribosomal peptide synthetases (NRPS) (12/15), while 3-347^T^ encoded for the production of terpenes. NRPSs and PKSs were enzyme complexes responsible for the formation of non-ribosomal peptides and polyketides, which formed two large and important groups of natural products with different chemical structures that yielded a wide range of biologically active compounds (34). Terpene biosynthetic clusters presented in both strains were all related to the synthesis of carotenoids, which could be the cause of the yellow colony coloration observed after three days of growth. Carotenoids were natural pigments with antioxidant properties (35). Notably, one carotenoid biosynthetic gene cluster in strain 1117^T^ exhibited 28.0 % similarity to that of *Algoriphagus* sp. KK10202C. Additionally, terpenoids, well-known natural products synthesized via the mevalonate pathway (36), were likely producible by these strains. In contrast to strain 1117^T^, strain 3-347^T^ had a specific RRE-containing cluster that may encode lanthipeptide. Lanthipeptides demonstrated diverse biological activities, including antibacterial (37), antiviral (38), suggesting that lanthipeptides produced by strain 3-347^T^ may play a protective role for red macroalgae.

### Metabolic pathways analysis

Secondly, genomic functional annotation revealed that strains 1117^T^ and 3-347^T^ possess 39 and 42 abundant and complete metabolic pathway modules, respectively. These modules encompass carbohydrate metabolism, energy metabolism, lipid metabolism, nucleotide metabolism, amino acid metabolism, glycan metabolism and metabolism of cofactors and vitamins. Furthermore, KEGG annotation of amino acid sequences assigned putative functions to 1,485 genes (39.3 %) in strain 1117^T^ and 1,533 genes (39.0 %) in 3-347^T^. The presence of these complete metabolic pathway modules and functional genes will also increase their competitiveness in colonizing the surface of macroalgae and facilitate their adaptation to the habitat. Fig. 2 presents heatmaps of metabolic pathways for all strains in the 16S rRNA phylogenetic tree, with detailed metabolic reconstruction of strains 1117^T^, 3-347^T^, and related strains of *Flavobacteriaceae* (Table S8). Next, we reconstructed the important metabolic pathways of strains 1117^T^ and 3-347^T^, including the complete heme biosynthesis metabolic pathway (Fig. S6A) and CMP-KDO biosynthesis metabolic pathway (Fig. S6B), which are critical for their colonization and survival on macroalgal surfaces. Heme, an essential cofactor, plays vital roles in respiration, gas sensing, and detoxification of reactive oxygen species (39). Heme biosynthesis pathways are present in most aerobic organisms across all three domains of life, and variations in these pathways among bacterial species offer potential targets for novel antimicrobial development (40). In addition, CMP-KDO synthetase [EC:2.7.7.38] was involved in the biosynthesis of lipopolysaccharide (LPS) which was an essential component of the outer membrane of gram-negative bacteria (41). The ability of strains 1117^T^ and 3-347^T^ to synthesize LPS enhances their extracellular membrane integrity, promoting growth and antibiotic resistance. Furthermore, the presence of TonB-dependent transporter genes in PUL-like regions of both strains suggests their potential to transport vitamin B12 to green and red macroalgae, respectively, supporting a symbiotic relationship. This aligns with previous findings that algae acquire vitamin B_12_ through bacterial symbiosis (42). These metabolic capabilities collectively enhance the ecological fitness of strains 1117^T^ and 3-347^T^, enabling them to thrive on macroalgal surfaces.

**Fig. 2.**
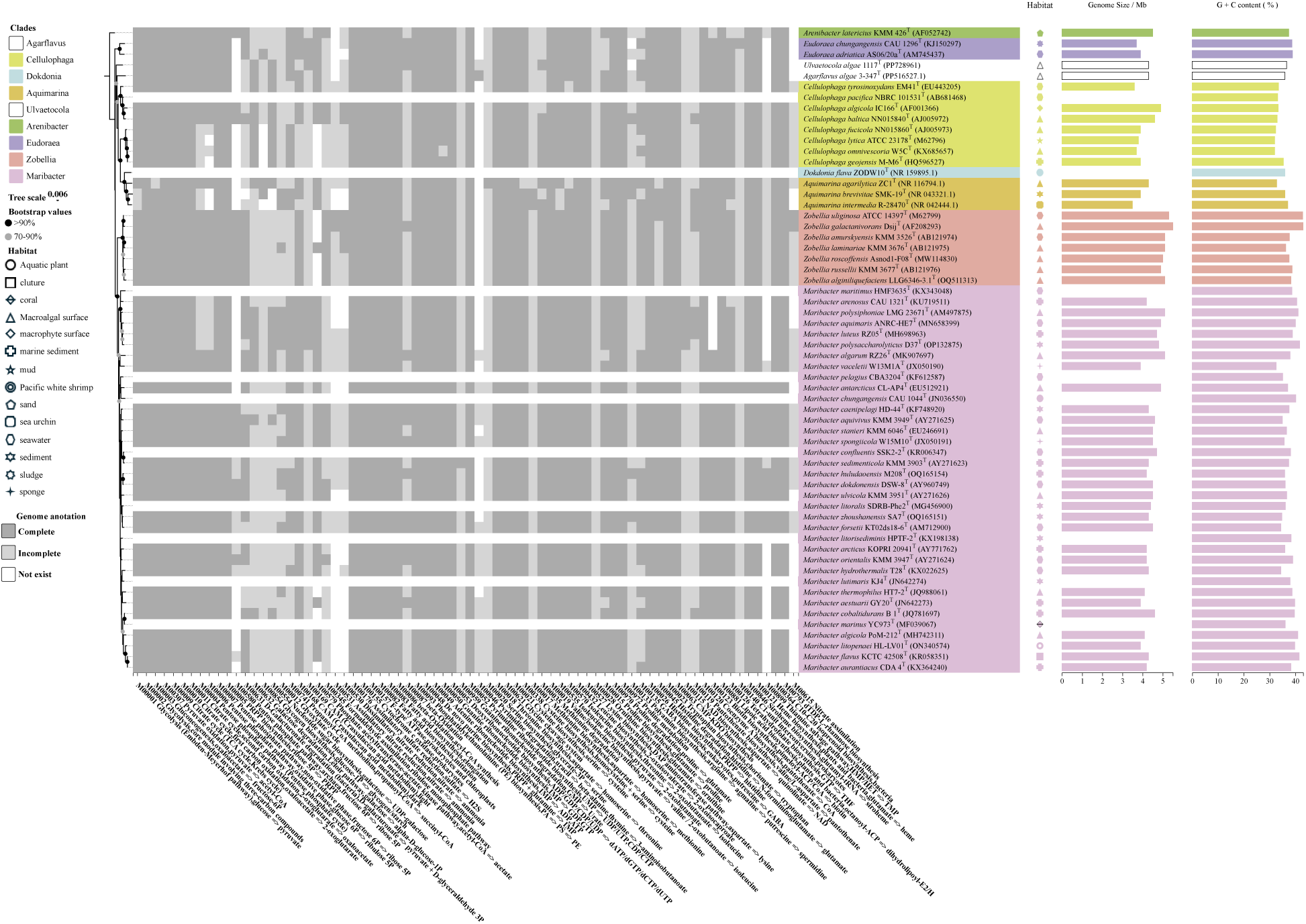
Taxonomic classification of strains 1117^T^ and 3-347^T^ based on the topological structure of 16s rRNA phylogenetic tree. *Ulvaetocola* and *Agarflavus* belong to different branches and positions and are two novel genera. Different colors are used to represent different clades. Black filled circles indicates that the bootstrap values are greater than 90.0 %, and gray filled circle indicates bootstrap values between 70.0-90.0 %. Bar, 0.006 substitutions per nucleotide position. Heat map shows metabolic pathways of different bacteria. The shade of the color indicates the integrity of the metabolic pathway, with dark grey indicating an intact metabolic pathway. In addition, different shapes, such as circles, squares and triangles, show the habitats of all the bacteria. Genome size (Mb) and G+C content (%) of different bacterial genera from different sources are represented by the bar graphs on the right.

Unlike 1117^T^, strain 3-347^T^ encoded glucose-1-phosphate thymidylyltransferase (EC:2.7.7.24), dTDP-glucose 4,6-dehydratase (EC:4.2.1.46), dTDP-4-dehydrorhamnose 3,5-epimerase (EC:5.1.3.13) and dTDP-4-dehydrorhamnose reductase (EC:1.1.1.133), suggesting that it may have the ability to biosynthesise dTDP-L rhamnose, as shown in Fig. S6C. Additionally, it had genes bioF (encoding 8-amino-7-oxononanoate synthase), bioA (encoding adenosylmethionine-8-amino-7-oxononanoate aminotransferase), bioD (encoding dethiobiotin synthetase) and bioB (encoding biotin synthase), demonstrating its ability to synthesize biotin (Fig. S6D). This suggested the existence of a potential symbiotic relationship between strain 3-347^T^ and red macroalgae (42), validating its isolation site. Specifically, strain 3-347^T^ may provide vitamin B_7_ (biotin) to macroalgae, supporting macroalgal growth and survival on their surfaces. This metabolic versatility likely contributes to the strain’s competitive advantage in colonizing and thriving in macroalgal habitats.

### Metagenomes and Habitat distribution analysis

Through genomic function analysis, we found that macroalgal epiphytic bacteria possess abilities to adapt and survive macroalgal surface environment. In order to determine their relative abundance in the samples, we compared the Reads per kilobase per million (RPKM) values of strains isolated from the surface of green and red macroalgae, including strains 1117^T^, 3-347^T^, four macroalgal epiphytic bacteria (2-919, 2-616, 2-621 and 2-787) and two *Ulvaetocola* MAGs (detailed values see Table S9). The results showed that the RPKM values of all strains and MAGs were much higher in macroalgae than in seawater and sediment (Fig. 3). This suggests that strains are highest in relative abundance in macroalgae, consistent with the fact that they are isolated from the surface of marine macroalgae. Strain 3-347^T^ had a significantly higher RPKM values in seawater than sediment, while the other strains were only slightly higher than in sediment. This suggested that strain 3-347^T^ was mainly distributed on the surface of the macroalgae and to a lesser extent in seawater. Strain 1117^T^ and other strains were mostly distributed on the macroalgal surface, but also in seawater and sediments. This suggests that strains, through their own metabolic and functional properties, exhibits greater competitiveness on the macroalgal surface and dominates the ecological competition.

**Fig. 3.**
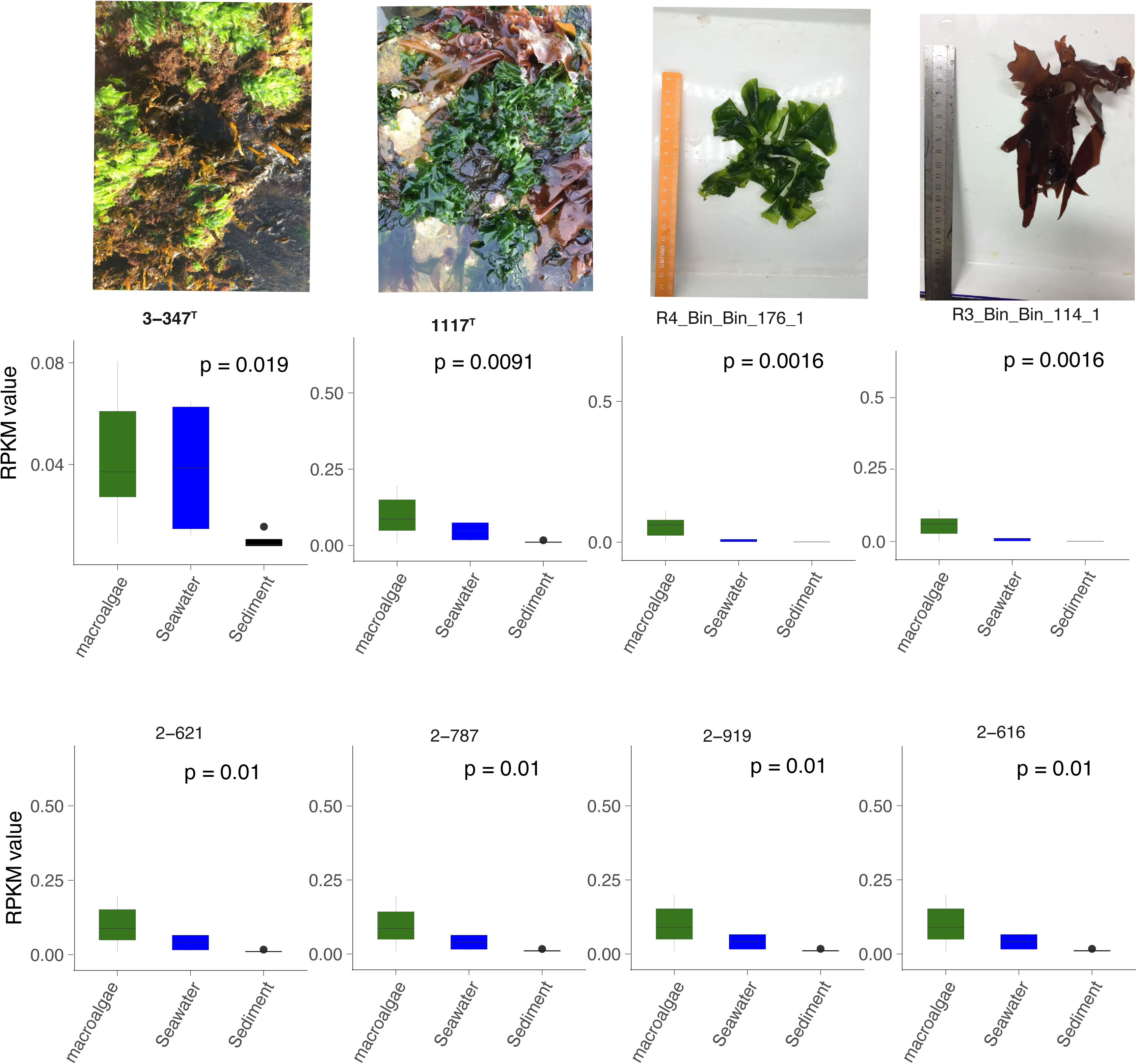
Photographs of the native environment from fall and spring sampling environments and two new genera of host marine macroalgae (Weihai, China). Metagenomes analysis of RPKM values of strains 1117^T^, 3-347^T^, four macroalgal epiphytic bacteria (2-919, 2-616, 2-621 and 2-787) and two *Ulvaetocola* MAGs. The RPKM values are based on read recruitment from sample metagenomes with a 97% identity threshold. Box plot results show that the RPKM values of all strains in macroalgae is much higher than in seawater and sediment, especially strains 1117^T^ and 3-347^T^. These strains are also distributed in seawater and sediment, suggesting a wider distribution.

However, due to the limitations in the number and types of samples, we utilized the Global Ecological Database (https://microbeatlas.org/mapseq) to validate the above conclusions (Fig. 4). Sequences with 97.0 % or more similarity to strains 1117^T^ and 3-347^T^ were found in 642 and 4,315 samples worldwide, respectively (detailed information see Table S10). These sequences were distributed across a wide range of environments, including marine habitats (e.g., biofilms, beaches, corals, sponges, marine sediments, seawater, and macroalgae), associated hosts (humans, animals, and plants), freshwater, and soils (Fig. 4). Strain 1117^T^ demonstrated a broad global distribution, with its relative abundance being highest in macroalgae and seawater, followed by biofilms and other marine environments (Fig. 4A). It is predominantly distributed in coastal seawater and on the macroalgal surface, with a minor presence in human-associated environments and soils. This distribution aligns with its initial isolation from the surface of green macroalgae. Metagenomes confirmed that strain 1117^T^ exhibits high abundance on macroalgal surfaces, as well as in other marine environments, such as seawater, sediments and corals. This is consistent with global ecological distribution studies. In contrast, strain 3-347^T^ was mostly distributed in marine sediment and marine, higher than the relative abundance in beach (Fig. 4B). These are still part of the marine habitat and our metagenomes results also confirm its high abundance in the marine environment, which would increase its competitive advantage on the macroalgal surface. Members of the genus *Ulvaetocola* and *Agarflavus* demonstrated a preference for marine habitats. Thus, data from global samples suggest that the widespread distribution of strains may be attributed to its metabolic diversity, environmental adaptability and interactions with macroalgae, may further expanding their range (Supplementary Materials).

**Fig. 4.**
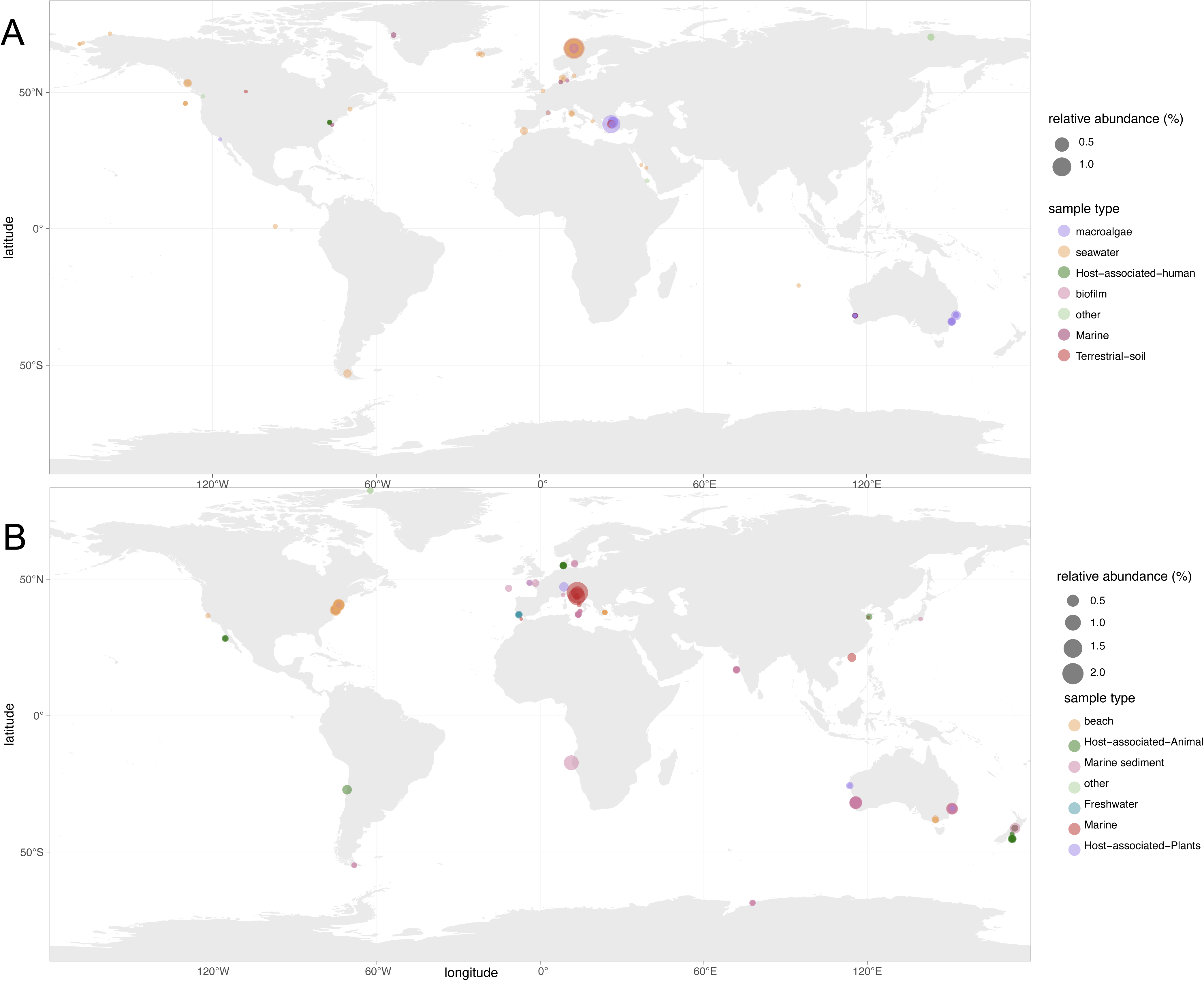
The worldwide environmental distribution of strains 1117^T^ and 3-347^T^ based on 16S rRNA gene sequences with a relative abundance above 0.01 %. The relative abundance of different samples types was estimated as a percentage of total readings calculated, with the size of the circle indicating high or low relative abundance, allowing for a more precise understanding of environmental distribution and preferences. (A) Strain 1117^T^was widely distributed globally, and its relative abundance was highest in seawater and macroalgae, followed by biofilm and marine. (B) Strain 3-347^T^ was mostly distributed in marine sediment and marine, higher than the relative abundance in beach.

### Prediction and annotation of CAZymes-rich gene clusters

In order to deeply investigate the polysaccharide degradation ability of macroalgal epiphytic bacteria, we selected strains 1117^T^ and 3-347^T^ for detailed analysis. We categorized the genes into four categories: (i) PUL-like clusters with CAZyme genes and an encoded TonB-dependent receptor, (ii) CAZyme-rich gene clusters (CGC) consisting solely of CAZymes, (iii) *susCD* loci without detectable CAZymes and (iv) PULs consisting of CAZyme genes and *susCD* pairs. Results showed strain 1117^T^ contained 5 PULs, 6 PUL-like clusters, 10 CGCs and 3 *susCD* pairs (Fig. 5A), while 3-347^T^ had 5 PULs, 8 PUL-like clusters, 16 CGCs and 2 *susCD* gene pairs (Fig. 5B). We then annotated their CAZyme genes and revealed that strain 1117^T^ had 163 CAZyme genes, including 50 GHs, 36 GTs, 31 CEs, 28 CBMs, 12 PLs and 6 AAs, while 3-347^T^ contained 177 CAZymes with 61 GHs, 44 GTs, 35 CEs, 24 CBMs, 7 PLs and 6 AAs. A list of CAZymes and its family activities of all bacteria in the genomic phylogenetic tree are shown in Tables S11, S12. Among these CAZymes, GHs were the greatest number of enzymes, which showed that they were more capable of degrading polysaccharides. Because polysaccharides were an important source of nutrients for storing energy and were conducive to microbial phycosphere colonization. Thus, macroalgal epiphytic bacteria, including 1117^T^ and 3-347^T^, can promote marine carbon cycling by degrading polysaccharides.

**Fig. 5.**
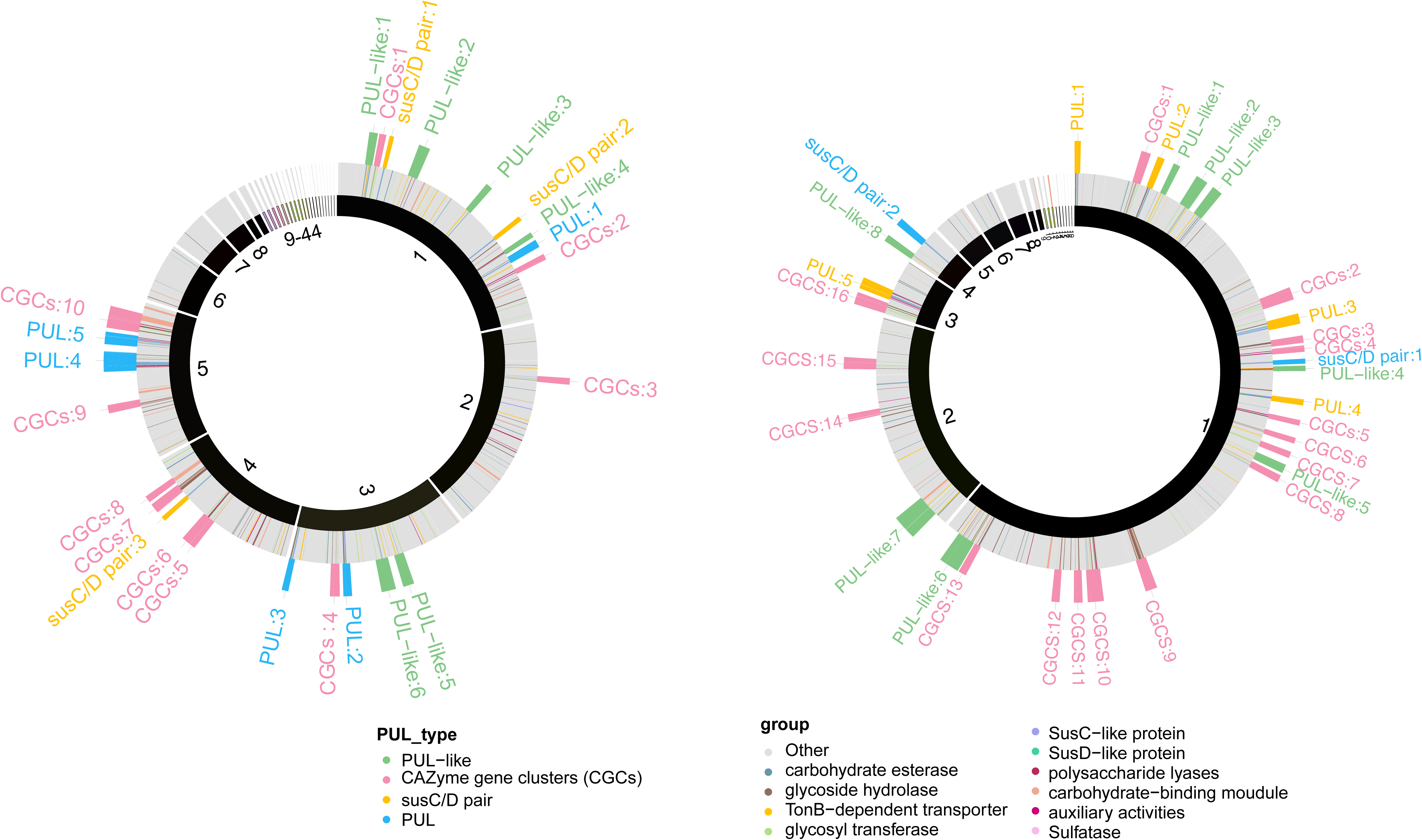
Distribution of different CAZymes groups and PUL_tpyes of strains 1117^T^ and 3-347^T^. They have different compositions and numbers of PUL_tpye, and thus have different ability to degrade different algal polysaccharides, demonstrating differences in polysaccharide degradation ability. Groups are diverse, found in all four PUL_tpyes and elsewhere, and their composition affects the polysaccharide degradation potential of strains. (A) Strain 1117^T^ contained 5 PULs, 6 PUL-like clusters, 10 CGCs and 3 *susCD* pairs. (B) Strain 3-347^T^ had 5 PULs, 8 PUL-like clusters, 16 CGCs and 2 *susCD* pairs.

### Putative substrate specificities

In the context of our study, macroalgal epiphytic bacteria exhibit significant potential for polysaccharide degradation, supported by the high prevalence and diversity of PULs in their genomes (15). This adaptation enhances their survival probability, facilitating more effective colonization and growth on algal surfaces. Consequently, we conducted extensive functional annotation of PULs in the obtained macroalgal epiphytic bacterial genomes. We revealed a wide range of as-yet-undescribed PULs in 299 genomes. A total of 610 annotated PULs can be linked to either dedicated polysaccharides or polysaccharide classes (Table S13), and some of the larger PULs were attributed to multiple polysaccharide substrates. Due to differences in PUL composition, PULs with common substrate predictions are not identical, but they can be systematically integrated into six different clusters. Of these, the highest number of PULs was annotated in Cluster5 (n=313), followed by Cluster2 (n=200), while the lowest number of PULs was annotated in Cluster3 (n=4). And clusters have their own polysaccharide degradation substrates, including alginates, xylan, starch, alpha-glucan, acetylxylan and xyloglucan substrates. Among 610 annotated PULs, forty-eight were specifically associated with acetylxylan degradation. Cluster1 comprised thirteen acetylxylan-targeting PULs (Fig. 6), bacterial taxa from diverse environmental sources (*Gelidium* sp., *Ulva* sp., and *Grateloupia* sp. from macroalgae and seawater) demonstrated acetylxylan degradation capabilities. These taxa exhibited conserved acetylxylan degradation patterns and similar gene compositions suggesting convergence of polysaccharide degradation potentials. Regarding alginate degradation, Cluster2 contained 200 of 610 annotated PULs. From these, nineteen representative PULs were selected for further analysis (Fig. 7). Bacterial isolates from marine macroalgae, sediment, and seawater environments demonstrated consistent alginate degradation potentials. However, some bacterial genomes lacked corresponding PULs, suggesting variability in polysaccharide degradation capacity.

**Fig. 6.**
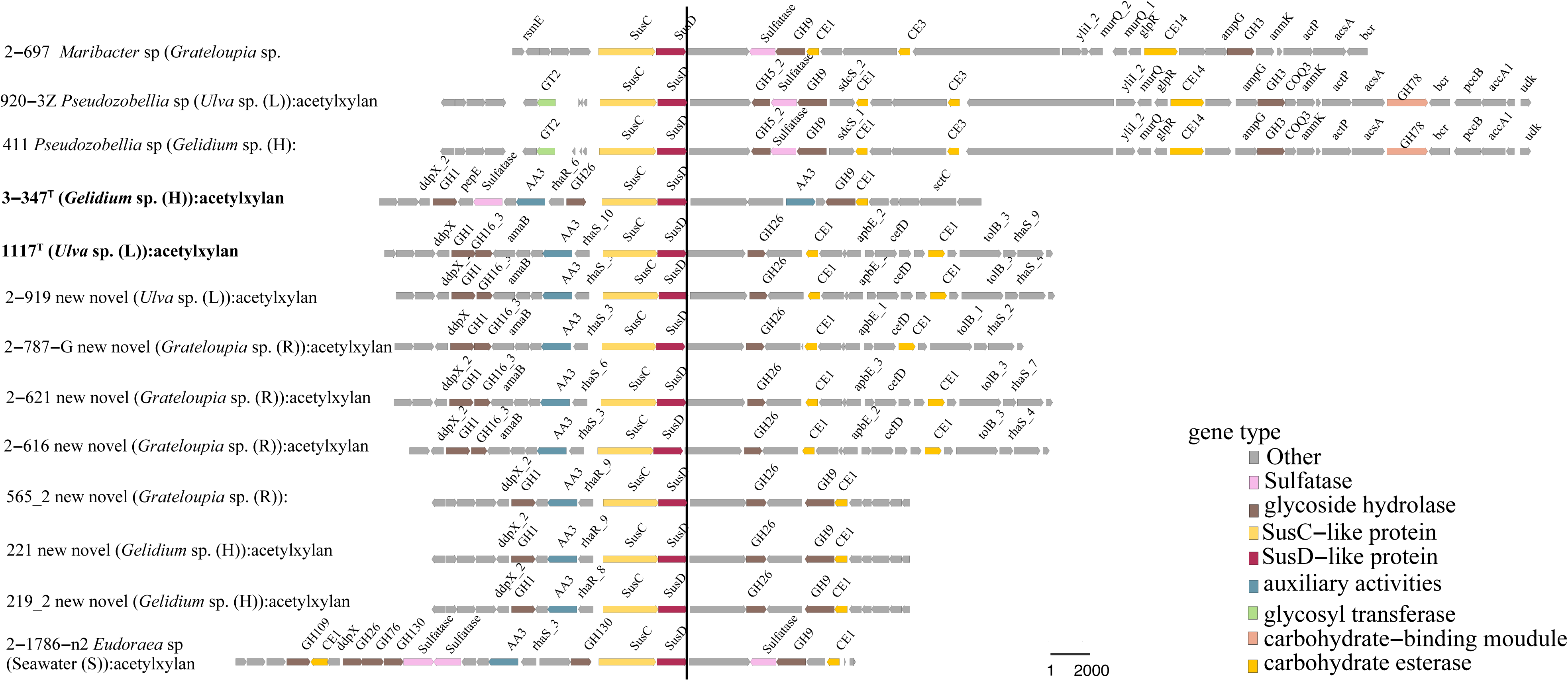
Cluster 1, associated with acetylxylan degradation, was identified in forty-eight out of 610 annotated PULs. Among these, thirteen representative PULs, including strains 1117^T^ and 3-347^T^, were analyzed. These PULs exhibited a consistent acetylxylan degradation pattern, with similar gene type composition and structural organization. Notably, bacteria from diverse taxa (e.g., *Gelidium* sp., *Ulva* sp. and *Grateloupia* sp.) and different environments (macroalgae and seawater) were found to possess identical PULs capable of degrading acetylxylan. A variety of gene types function synergistically to hydrolyze polysaccharide linkages. Additionally, other genes within these PULs serve specific functions, including those regulating bacterial growth and development, as well as those involved in polysaccharide transport.

**Fig. 7.**
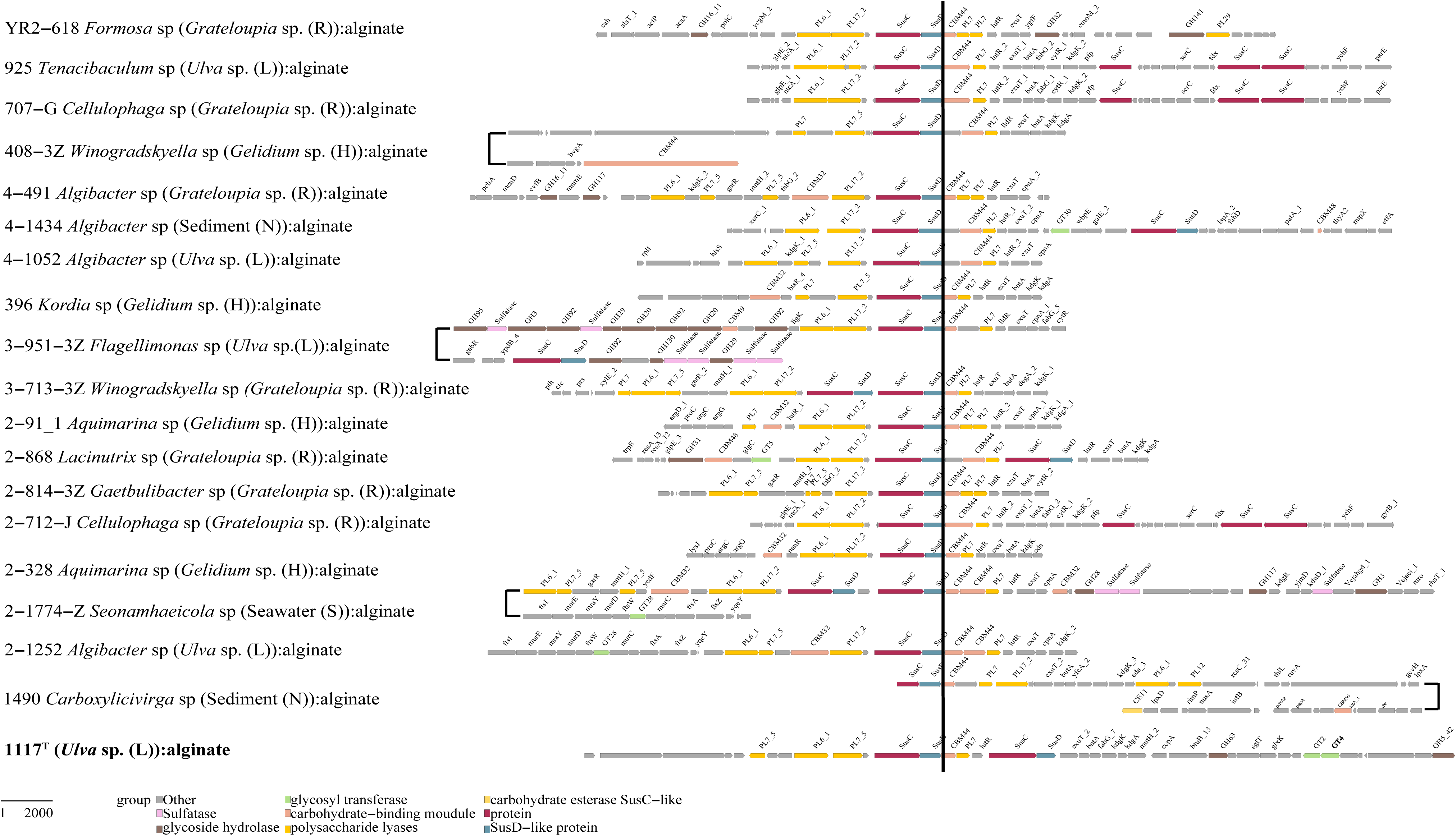
Nineteen out of the 200 annotated PULs associated with alginate degradation were selected for analysis, including strain 1117^T^. Compared to other clusters, Cluster2 exhibits the greatest length, structural complexity and genetic diversity. Bacteria originating from marine macroalgae, sediment, and seawater were all found to degrade alginate. The diverse genotypes and complex organization of bacterial PULs within this cluster determine their polysaccharide degradation capabilities. Additionally, other genes within these PULs serve specific functions, including those regulating bacterial growth and development, as well as those involved in polysaccharide transport.

These data revealed that epiphytic bacteria derived from different macroalgae and isolation sources possess similar PULs for degrading identical polysaccharides (Figs. 6, 7, S7, S8). For example, both 1117^T^ and 3-347^T^ are involved in acetylxylan and xyloglucan degradation (Figs. 6, S8). Certain bacteria may specialize in degrading one or a few specific polysaccharides, while others exhibit a broader degradative capacity (Figs. 6, 7, S7, S8). For instance, strain 1117^T^ may degrade alginate and α-glucan (Figs. 7, S7C), while 3-347^T^ may degrade galactomannan (Fig. S7A).

### Xylan, acetylxylan and xyloglucan

Xylan, comprising linear chains of β-1,4-linked xylosyl residues, represents the second most abundant polysaccharide in plant biomass (43). It is structurally complex and its degradation requires a variety of CAZyme gene families, including GH10, 11, 30, 43, 51, 67, 115, and the CE1, 4 families (44). Of GH family, GH10 was first reported (45). Genomic analysis revealed that xylan-targeting PULs in strains contained GH30_1 and GH74 genes. Notably, strain 3-347^T^ additionally possessed genes from the GH43_11 family. Both strains encoded PULs containing CE families CE1, 3, 7, which were potentially involved in acetyl group removal. This genetic architecture suggested that these PULs may target oligosaccharides containing acetylated xylose residues, indicating the presence of xylan acetylation in marine macroalgae. Furthermore, extensive PUL analysis identified multiple CE families, with 1117^T^ containing CE7 and 3-347^T^ possessing CE17. These enzyme families function as glucuronyl esterases and acetylxylan esterases, respectively, playing crucial roles in lignocellulose degradation and acetyl group removal from hemicelluloses (46).

The side chain of xylan had an acetyl modification, which was attached to the C2 or C3 position of the xylose residue via an ester bond, giving acetylxylan (44). Forty-eight bacterial strains harboring Cluster1, characterized by diverse CAZyme genes composition including sulfatase genes (Fig. 6), exhibited specific acetylxylan degradation capabilities. Phylogenetically distinct bacterial taxa (*Gelidium* sp., *Ulva* sp. and *Grateloupia* sp.) isolated from various ecological niches—including green macroalgae, two distinct red macroalgae species, and seawater environments—all possessed acetylxylan-targeting PULs, indicating that strains had evolved with more generalized polysaccharide degradation potential for growth enhancement. Similarly, Cluster4 represented an alternative genetic configuration for acetylxylan degradation, although its CAZyme gene composition was different from Cluster1.

Furthermore, Cluster6 demonstrated xyloglucan degradation capacity (Fig. S8). Genomic analysis revealed that this cluster was present in bacterial strains isolated from marine green macroalgae and red macroalgae. This phylogenetic distribution suggested that xyloglucan degradation potential represents a conserved functional trait among macroalgal epiphytic bacteria (47). It had been shown that fungal esterase and xylanase activities act synergistically in hydrolyzing acetylxylan (48). Their genome had genes for fungal family (DUF1776), beta-1,3-xylanase function, xylulose kinase and other function. These enzymes hydrolyzed acetylxylan into xylose, xylobiose, and small amounts of other monosaccharides. Xylose was first phosphorylated to xylulose-5-phosphate by xylulose kinase and then converted to glucose-6-phosphate via the pentose phosphate pathway. It thus sequentially entered the glycolysis and TCA cycles, generating carbon dioxide and producing large amounts of ATP. This suggested that strains possessing the appropriate enzymes could degrade acetylxylan to satisfy their nutritional and energy needs.

### Alginates

Alginates were linear co-polymers consisting of homopolymeric blocks of (1→4)-linked β-D-mannuronate and α-L-guluronate residue (49). Both it and its degradation product (alginate oligosaccharides) were rich in biological activities such as antioxidant activity and antimicrobial activity, which played an important role in the marine carbon cycle (50). PL7 family was first reported in 1982 (51), PL5 family in 1987 (52) and PL15 in 2000 (53). They all contained alginate lyase and were mainly distributed in marine bacteria (54). PL6_1, 7, 7_5 and 15_1 family alginate lyases of strain 1117^T^ could degrade alginate, suggesting that it could degrade alginates, which was consistent with the experimentally verified results (see Supplementary Materials). Bacterial isolated obtained from diverse marine environments—including green macroalgae, two distinct red macroalgae, seawater, and marine sediments—were found to possess alginate-targeting PULs (Fig. 7). Nineteen isolates harbor PULs rich in alginate-targeting CAZyme genes, e.g., from families GH1, 26, 9, 3, 16_3 and 125. In comparison to other clusters, Cluster2 exhibited the most complex genetic architecture and enriched with diverse CAZyme genes, particularly GH family genes. The genetic diversity observed in these PULs directly correlates with their polysaccharide degradation capabilities. The ubiquitous distribution of alginate-degrading bacteria across various marine habitats suggested that alginate degradation represented a conserved functional trait among macroalgal epiphytic bacteria.

Additionally, alginate was broken down into β-D-mannuronate and α-L-guluronate by the action of alginate lyase. And the glyoxylate pathway was accomplished through a series of enzymatic reactions. The key step in it involved the conversion of the two glyoxylates to UDP-mannose and UDP-gulose, respectively, which were then released as glucose by sugar unit transfer or hydrolysis reactions. Thus, glucuronic acid was converted to glucose, which entered the glycolysis (Embden-Meyerhof pathway) and was converted to pyruvate and produced large amounts of energy. Pyruvate entered the mitochondria and goes through the TCA cycle and oxidative phosphorylation process to produce carbon dioxide and more ATP molecules. Strain 1117^T^ had complete glycolysis and tricarboxylic acid cycle pathways to fulfill energy and metabolic requirements (Table S8).

### Alpha-glucan

Starch (α-1,4- and α-1,6-glucan) was widely found in terrestrial plants, macroalgae and animals, and thus played a central role in life as a principal store of chemical energy (55) (56). GH13, GH15, GH31, GH57, GH65 and GH77 family enzymes were involved in the degradation of alpha-glucan (57, 58), with GH13 being the first reported (59). This suggested that strain 1117^T^ and 3-347^T^ hydrolyzed starch, veryfing the results of the physiological experiments (see Supplementary Materials). GH65 and GH13_38 enzymes genes in PUL2 of strain 1117^T^ involved in structurally simple and non-branched α-glucan degradation, like starch. Furthermore, Cluster5 was identified as specifically targeting α-glucan degradation. This cluster comprised 303 bacterial PULs derived from diverse marine environments, including green macroalgae, two distinct red macroalgae species, seawater, and marine sediments (Fig. S7C). The cluster exhibited structural homogeneity, a relatively compact genomic architecture, and an enrichment of GH genes, particularly GH13_38 and GH65. In terms of strain 3-347^T^, 1 GH13_38 that encoded maltose-1-phosphate maltosyltransferase and 1 GH65 that encoded Maltose phosphorylase were found in PUL2, which represented the PULs for α-glucans.

Furthermore, we identified a large number of CAZyme genes outside the four main functional categories. These genes potentially encode enzymes capable of degrading diverse polysaccharide substrates, including but not limited to laminarins, pectin, chitin, peptidoglycan, agars, N-glycans, β-mannans, trehalose, porphyrans, carrageenans, amylosucrase and galactomannan. Comprehensive analyses of these polysaccharide degradation pathways and detailed characterization of GHs, GTs, AAs and CEs family genes are provided in Supplementary Materials.

### Growth curve of polysaccharide substrates

In order to validate the predicted conclusions about polysaccharide substrates for macroalgal epiphytic bacteria, we selected four common laboratory polysaccharides for growth curve experiments on strains 1117^T^ and 3-347^T^. We compared their growth rates in media containing starch, xylose, β-1,3-glucan and carboxymethylcellulose sodium. We subtracted the initial optical density value so that the growth curves start at zero. And we set up a control group, recorded detailed data (Table S14) and plotted growth curves (Fig. S9). Strain 1117^T^ grew better in the media which contain carboxymethylcellulose sodium and β-1,3-glucan separately compared to control group, while 3-347^T^ grew better in the media which contains starch, β-1,3-glucan, and carboxymethylcellulose sodium separately. The experimentally verified results of strains 1117^T^ and 3-347^T^ for the degradation of starch, β-1,3-glucan and carboxymethylcellulose sodium were consistent with the results of annotated PULs. However, due to the wide variety of xylose types, they were not able to degrade the xylan of the validation experiment, so the growth rate in the growth curve was lower than control group.

## Discussion

The surfaces of marine macroalgae are enriched with specific microbial taxa, which have co-evolved with their hosts over extended evolutionary timescales (17). The mutual selection between bacteria and their hosts is a critical driver shaping the composition of epiphytic microbial communities (60, 61). Studies of these two novel genera have elucidated that macroalgal epiphytic bacteria may acquire their ecological niches and competitive advantages by synthesizing secondary metabolites. Strains 1117^T^ and 3-347^T^ not only act as carotenoid lactobacilli for the production of carotenoids, but also possess abundant BGCs (Fig. S5), which have a great potential for the production of diverse biologically active compounds, which are essential for maintaining community composition (62). This suggests they occupy specific ecological niches on macroalgal surfaces, enhancing their competitiveness for colonization and survival. We also hypothesize that they may have potential evolutionary relationships for better environmental adaptation. Our findings reveal that the surface of marine macroalgae is rich in bacteria, encompassing numerous previously undescribed novel genera and species within the phylum *Bacteroidota* (Fig. 1). Through comprehensive taxonomic classification, genomic sequencing, physiological and chemotaxonomic analysis, we identified and established two novel genera originating from the surfaces of marine macroalgae (detailed description see Supplementary Materials). Genomic function analysis of two new genera revealed more complete metabolic pathways (Fig. 2; Table S8), high relative abundance on the macroalgal surface, strong ability to synthesize secondary metabolites (Fig. S5) and abundant PULs (Fig. 5). We also conducted metagenomes (Fig. 3; Table S9) and global ecological distribution analyses (Fig. 4; Table S10). These abilities and properties help them to better colonize, grow and adapt to specific environments on the macroalgal surface.

We analyzed the proportions of novel species and genera within the phylum *Bacteroidota* across different macroalgae samples (Fig. 1; Table S1). We found that the highest percentages of novel species and genera were present in *Grateloupia* sp. samples, while the lowest percentages were observed in *Saccharina* sp. samples. Furthermore, we conducted metagenomes analysis of the RPKM values of some *Bacteroidota* in four macroalgae samples (Fig. 3; Table S9). The results revealed that the relative abundance of *Bacteroidota*, as well as the average relative abundance, was highest in *Ulva* sp. The RPKM values of macroalgal epiphytic bacteria were significantly higher in macroalgae samples compared to those in seawater and sediment.

The study revealed that macroalgal epiphytic bacteria possess the capability to degrade polysaccharides, targeting both similar and distinct algal polysaccharides (Figs. 6, 7). However, this capability does not exhibit a clear correlation with taxonomic classification. These epiphytic bacteria harbor a diverse and complex array of PULs, indicating their potential for broadly degrading algal polysaccharides (63). Annotation of 299 macroalgal epiphytic bacterial genomes demonstrated that bacteria from different genera and habitats share identical PULs capable of degrading the same algal polysaccharides (Table S13). This finding supports the hypothesis of horizontal gene transfer (HGT) facilitating the exchange of PULs among members of family *Flavobacteriaceae* (64). For example, strains 1117^T^ and 3-347^T^ belong to two different novel genera from green and red macroalgae, respectively, but both of them had Cluster1 to degrade acetylxylan and Cluster6 to degrade xyloglucan (Figs. 6, S8). This convergence in PUL composition is attributed to the core microbial communities on macroalgal surfaces, which have evolved PULs capable of degrading diverse macroalgal polysaccharides as an adaptive strategy for colonization across various macroalgal species (65). Furthermore, bacteria of the same genus differ in their PULs, which enables them to degrade different algal polysaccharides. This finding supports the hypothesis that the functional roles of PULs in a species are more strongly influenced by its unique ecological niche than by its phylogenetic lineage (49). Macroalgal epiphytic bacteria from different genera and habitats were found to possess distinct functional PULs. The variability in bacterial polysaccharide-degrading capabilities reflects their specificity and diversity in metabolizing polysaccharide substrates, which is closely linked to their growth, development, colonization and environmental adaptation (65). This functional differentiation enables bacteria to occupy distinct ecological niches (63), reducing direct competition among them and thereby contributing to the stability of microbial communities (66). For instance, strain 1117^T^ harbors Cluster2 and Cluster5 for degrading alginate and α-glucan (Figs. 7, S7C), while 3-347^T^ possesses Cluster3 for degrading galactomannan (Fig. S7A). These distinct PUL profiles enable two strains to degrade different polysaccharides, facilitating their adaptation to epiphytic lifestyles on green and red macroalgae.

Among the 15,324 genes identified in 610 annotated PULs, those targeting α-glucan constituted the largest proportion (42.2%), followed by those targeting alginate (32.9%). PULs associated with α-glucan degradation accounted for 39.8% (243/610) of all PULs in the isolates, while those targeting alginate represented 32.9% (201/610) (Table S13). The widespread prevalence of these degradation capabilities is likely associated with macroalgae, although they are also frequently observed in numerous marine isolates (49). This suggests that these polysaccharides are abundant substrates in isolated habitats, utilized by numerous bacteria, which may compete for these resources to survive (67). We further hypothesize that macroalgal epiphytic bacteria have evolved enhanced polysaccharide degradation potential (63). Diverse bacteria from various sources have developed the ability to degrade the most abundant polysaccharides in their respective habitats, thereby facilitating their growth and survival (65). Macroalgae epiphytic bacteria utilized algal polysaccharides as energy and nutrients, which participated in the marine carbon cycle (68) and played an important role in the marine ecosystem (17, 69).

## Materials and methods

### Bacterial isolation and cultivation

For strain isolation, whole macroalgae were cut into 9 cm^2^ pieces and rinsed three times with sterile seawater. Then 10 g of these pieces were washed with 10 ml of sterile seawater (rotary shaker, 170 rpm, 30 min, 25°C). One milliliter aliquots were diluted gradually with sterile seawater to 10^-6^, then 100 μl were spread and incubated with a standard dilutionplate method in the modified marine agar 2216 medium: 18 g sea salt (Sigma-Aldrich, St. Louis, MO, USA), 1.5 g peptone, 0.3 g yeast extract, 0.3 g sodium pyruvate, 0.3 g glucose, 15 g agar, 2 g alginate, 2 g starch, 2 g carrageenan, 2 g cellulose and 0.5 mg vitamin B_12_ (28). After 21 days of incubation at 28.0°C, single colonies were selected, purified and identified, which were cultivated in MA and cryopreserved containing 20.0 % (v/v) glycerol at -80.0°C. Light microscopy (E600, Nikon) and scanning electron microscopy (Nova NanoSEM450, FEI) were used to access the size and cell morphology.

### 16S rRNA gene sequence amplify

The PCR with universal primers 27F (5’-AGAGTTTGATCCTGGCTCAG-3’) and 1492R (5’-TACGGYTACCTTGTTACGACTT-3’) (70) was used to amplify the partial 16S rRNA genes of the colonies grown on MA. The complete 16S rRNA gene sequences of strains 1117^T^ and 3-347^T^ were extracted from the draff genomes by ContEst16S (https://www.ezbiocloud.net/tools/contest16s). For comparative analysis, the determined 16S rRNA gene sequence was submitted to the National Centre for Biotechnology Information (NCBI) GenBank database. The similarities of the 16S rRNA gene between the strain and related phylogenetic neighbors were identified and calculated in NCBI blast system (https://www.ncbi.nlm.nih.gov/) and EzBioCloud Database (71). Phylogenetic analysis was performed using MEGA X software (http://www.megasoftware.net/) (72). Phylogenetic trees were reconstructed using the Neighbor-Joining (NJ) (73), Maximum-Likelihood (ML) (74), and Minimum-Evolution (ME) (75). The genetic distances were calculated by the Kimura two-parameter model (76), and the topology was estimated by bootstrap analyses based on 1000 replications for each method.

### Selection of phenotypes for delimiting genera

For each species, the original isolation paper was used to retrieve phenotypic data. If data were missing for specific characters, then other sources were used. Several characters were collected for all species such as Gram type, morphology, motility, flagellation, growth conditions, reduction of nitrate, catalase activity, oxidase activity, DNA G+C content, isolation habitat, and results from the API-(ZYM, 20E) system. Individual characters were identified that appeared to vary with the core-genome phylogeny and were then used to inform taxonomic assignments.

### Genome extraction, sequencing, assembly, binning and annotation

Genomic DNA was extracted using a bacterial genomic DNA extraction kit (Takara, http://www.takara.com.cn, China). Quality was checked by running the samples on 1 % sodium boric acid agarose gels and measuring DNA concentrations using a NanoDrop 1000 spectrophotometer (Thermo Fisher Scientific, Waltham, MA, USA) based on 260/280 nm and 260/230 nm absorbance ratios. The draft genome sequencing was carried out by Beijing Novogene Biotechnology (Beijing, China) on the Illumina NovaSeq 6000 platform (Illumina, San Diego, CA, USA) with 150 bp PE reads at ≥100×coverage. Raw sequencing data were generated using Illumina base-calling software CASAVA v1.8.2^4^, and the high-quality paired-end reads were assembled using SPAdes v3.9.1 (–careful –covcutoff) with k-mer sizes from 27 to 127 bp and a minimum scaffold length of 200 bp (28). The genome were predicted using Prodigal v2.6.3 (77) and annotated with Prokka (78). Genes involved in metabolic pathways were analyzed in detail using information from the KEGG’s KofamKOALA server (79) (https://www.kegg.jp/blastkoala/) and the Rapid Annotations using Subsystem Technology (RAST) server (80). The antiSMASH 7.0 database was used to predict the presence of gene clusters encoding secondary metabolites (81).

The G+C content of the chromosomal DNA was calculated using genome sequence. The average nucleotide identity (ANI) was measured using the OAT software (82). And the digital DNA-DNA hybridization (dDDH) values were obtained using the genome-to-genome distance calculator (GGDC) website (83). The percentage of conserved proteins (POCP) (A proposed genus boundary for the prokaryotes based on genomic insights) was used to delineate genera and estimated their evolutionary. Studies had shown that POCP could be used as a genomic index for prokaryotic genus delimitation (30).

### Metagenome-assembled genomes (MAGs), Taxonomic inference of MAGs and abundance analysis

A total of 1.4 Tbp (avg. 65 Gbp per metagenome) were generated (28). Read quality filtering was done with BBDuk v35.14 (http://bbtools.jgi.doe.gov) and verified with FastQC v0.11. Reads from each sample were subsequently assembled individually using MEGAHIT v1.2.9 (84) with a minimum scaffold length of 2.5 kbp. BAM files were generated for each metagenome by mapping reads onto assemblies with BBMap v38.86 (minid = 0.99, idfilter = 0.97, fast = t and nodisk = t.) Initial binning was performed from within anvi’o v6.2 (85) using CONCOCT v0.4.0 (86), MaxBin v2.1.1 (87) and MetaBAT v0.2 (88). Resulting bins were combined with DAS Tool v1.1 (89) in order to find an optimal set. MAGs were denoted by an initial capital letter specifying the sample (B = *Saccharina*, L = *Ulva*, H = *Grateloupia*, R = *Gelidium*, S = seawater, N = sediment), followed by a number representing the season (1 = autumn, 2 = winter, 3 = spring, 4 = summer), followed by the binning program, and a terminal numeric identifier. In order to determine MAGs and genomes abundance, metagenomic raw reads were mapped to MAGs using BBMap (minid=99). RPKM was calculated based on MAG length and number of reads mapped.

### Habitat distribution analysis

The 16S high-throughput sequencing data for millions of global samples from multiple habitats were acquired from the Microbial Atlas Project (90) (MAP, https://microbeatlas.org/, accessed 30 July 2024). Each operational taxonomic unit (OTU) was classified by vsearch (91) using the silva SSU Ref NR 99 138.1 dataset. OTUs that displayed more than 97.0 % 16S rRNA gene sequence similarity to strain 1117^T^ and 3-347^T^ were identified in the genus *Ulvaetocola* and *Agarflavus*. The habitat distribution of genus *Ulvaetocola* and *Agarflavus* were determined through studying the bacterial abundance of the genus *Ulvaetocola* and *Agarflavus* in samples from all over the world.

### Prediction and annotation of PULs and CAZymes-rich gene clusters

The prediction and annotation of CAZymes-rich and PULs gene clusters were conducted following the methodology proposed by Dechen et al (28) and R (https://www.r-project.org) generated the related figure. As of July 2020, genes encoding CAZymes were annotated according to Krüger et al (92) description using a combination of HMMER searches against the dbCAN v2.0.11 (93) database and Diamond v0.9.24.125 searches against the CAZy database (94). The related HMMER and TIGRFAM profiles were used to predict the genes encoding sulfatases, *SusC*-, and *SusD*-like proteins. A sliding window of ten genes was used to predict PULs and other CAZyme-rich gene clusters, as stated in Francis et al (95). Additionally, we used dbCAN2 to find these clusters (93).

## Acknowledgements

This work was supported by the Science Foundation for Youths of Shandong Province (ZR2023QC197), the Science and Technology Fundamental Resources Investigation Program (Grant number 2022FY101100) and the National Natural Science Foundation of China (92351301). The scanning electron microscopy was supported by the Physical-Chemical Materials Analytical and Testing Center of Shandong University at Weihai.

## Data availability

The GenBank accession number for the 16S rRNA gene sequence of strains 1117^T^ and 3-347^T^ is PP728961.1 and PP516527.1, respectively. The draft genome of strains 1117^T^ and 3-347^T^ had been deposited in GenBank under the accession number CANMCC000000000 and CANKZG000000000, respectively. BioSample: SAMEA112156868, SAMEA112156860; BioProject: PRJEB57783. Sequences are available from the European Nucleotide Archive under accessions PRJEB50838 (metagenomes and MAGs), and PRJEB57783 (genomes of cultured bacteria).

## Declarations

Conflicts of interest Authors declare that there was no conflict of interest. This article does not contain any studies with animals performed by any of the authors.

## Ethics approval

Not applicable.

## Consent to participate

Informed consent was obtained from all individual participants included in the study.

## Consent for publication

All authors have agreed to it being published.

### Abbreviations

KCTC: Korea Collection for Type Culture
MCCC: Marine Culture Collection of China
MA: marine agar
NCBI: National Centre for Biotechnology Information
RPKM: Reads per kilobase per million
ANI: average nucleotide identity
dDDH: digital DNA–DNA hybridization
POCP: The percentage of conserved proteins
GGDC: Genome-to-Genome Distance Calculator
NJ: Neighbor-Joining
ML: Maximum-Likelihood
ME: Minimum-Evolution
PE: phosphatidylethanolamine
L: unidentified lipids
AL: unidentified aminolipid
PG: phosphatidylglycerol
MK-6: Menaquinone 6
GH: glycoside hydrolases
GT: glycosyl transferases
CE: carbohydrate esterases
CBM: carbohydrate-binding modules
AA: auxiliary activities
PL: polysaccharide lyases
CAZyme: carbohydrate-active enzyme
PUL: polysaccharide utilization loci
CGC: CAZyme-rich gene clusters
BGC: biosynthetic gene clusters
NRPS: non-ribosomal peptide synthetases.

